# The Backpack Quotient Filter: a dynamic and space-efficient data structure for querying *k*-mers with abundance

**DOI:** 10.1101/2024.02.15.580441

**Authors:** Victor Levallois, Francesco Andreace, Bertrand Le Gal, Yoann Dufresne, Pierre Peterlongo

## Abstract

Genomic data sequencing has become indispensable for elucidating the complexities of biological systems. As databases storing genomic information, such as the European Nucleotide Archive, continue to grow exponentially, efficient solutions for data manipulation are imperative. One fundamental operation that remains challenging is querying these databases to determine the presence or absence of specific sequences and their abundance within datasets.

This paper introduces a novel data structure indexing *k*-mers (substrings of length *k*), the Back-pack Quotient Filter (BQF), which serves as an alternative to the Counting Quotient Filter (CQF). The BQF offers enhanced space efficiency compared to the CQF while retaining key properties, including abundance information and dynamicity, with a negligible false positive rate, below 10^*−*5^%. The approach involves a redefinition of how abundance information is handled within the structure, alongside with an independent strategy for space efficiency.

We show that the BQF uses 4x less space than the CQF on some of the most complex data to index: sea-water metagenomics sequences. Furthermore, we show that space efficiency increases as the amount of data to be indexed increases, which is in line with the original objective of scaling to ever-larger datasets.

**Availability:** https://github.com/vicLeva/bqf

## 1 Introduction

Genomic data sequencing is a wonderful tool for understanding the ins and outs of biological systems. Sequencing produces numeric plain text, organized as reads in files. Most of these files are gathered in public databases like the European Nucleotidic Archive (ENA) [7] that weighs 54.5PB by early 2024. The size of the databases follows exponential growth, and thus we need appropriate solutions to manipulate the data it contains. One simple operation that we are not yet able to achieve is to query the database and then, for each dataset, answer if a sequence is present or absent. Even better, answer for each dataset how many times a sequence is present: its abundance. To this end, we use indexing data structures that can handle another representation of the data, making it easier to query afterward. Some of the current indexing data structures use sets of *k*-mers (substrings of length *k, k* usually in [20; 50]) as the representation to query. In this way, the proportion of shared *k*-mers between a query sequence and a dataset. The main operation is thus to determine for each *k*-mer in which indexed dataset it occurs and with what abundance (how many times it occurs in a dataset).

Due to the scale of databases to index, recognized tools often sacrifice precision for the sake of performance. This can be done through pseudo-alignment as defined in [2], breaking down the queried sequences into *k*-mers and comparing them against *k*-mers of the datasets, often organised in “colored de Bruijn graph” representation of as in Bifrost [12] or GGCAT [9]. Here, the graph construction is the main limitation of the methods. Other tools allow false-positive results by using Approximate Membership Queries (AMQ) data structures to enhance space efficiency [6, 4, 20, 14, 27, 16]. They all use trade-offs between size and false-positive rate. By taking advantage of DNA and *k*-mers properties (small alphabet, redundancy of consecutive *k*-mers), the use of a simple associative array with super-*k*-mers [15] whose minimisers [25] have been hashed with a minimal perfect hash function [23] can create exact and space efficient indexes such as SSHash [21, 22]. However, apart from being static, this method suffers from significant computational requirements and is considerably slower compared to the approach we suggest. Data structures form the core of the tools mentioned above. The choice of the structure impacts the performance and the range of operations available to the user. To illustrate, a Bloom filter [5] can insert elements after it has been built in memory, while an XOR filter [10, 11] has better space usage, but is static. A Quotient Filter [3] allows more dynamicity than a Bloom filter as it can enumerate inserted elements and thus relocate elements in a smaller or larger structure as needed. The Quotient Filter is the backbone of the Counting Quotient Filter (CQF) [19], which can retrieve not only the presence or absence of a *k*-mer, but also its abundance. However, this structure has the disadvantage of requiring a lot of space.

In this paper, we propose a new genomic data indexing structure, an alternative to the CQF called the “Backpack Quotient Filter” (BQF). It is more space-efficient than the CQF while still offering the same properties (abundance, dynamicity), at the cost of a negligible false-positive rate. We propose a novel way to handle the abundance information. We leave the trade-off choice between (total) space and counts encoding precision to the user. In addition, we use the Fimpera [26] scheme to reduce each element’s space usage. In total, our tests show that at the price of a false-positive rate below 10^*−*5^%, the BQF can index billions of elements and their abundance, using between 13 and 26 bits per element. Compared to existing solutions, the BQF has the fastest average query time, while being fully dynamic. It is, to our knowledge, the only data structure that cumulates these features.

## 2 Material and Methods

### 2.1 Preliminaries

#### 2.1.1 *k***-mers, pseudo-alignment, and indexing**

A *k*-mer is any sequence of given size *k*. It can be of any character but in our context a *k*-mer is a substring of a genomic sequence, *i.e*. made up of nucleotides (A,C,G,T). *k*-mers are to genomics what words are to natural language: this way we can compare sequences by comparing their words. The number of *k*-mers existing in two sequences provides a metric to measure the similarity between them, leading to the so called pseudo-alignment [2]. In order to efficiently perform pseudo-alignments between any queried sequence and a dataset, we index its *k*-mers. Doing so, it is possible to know in constant and fast time (see Results) whether a *k*-mer belongs to the dataset or not. Then, when querying a sequence *S*, all of its *k*-mers are individually queried in the index, enabling to compute the pseudo-alignment between *S* and each dataset of the databank.

In this article, the examples will use 32-mers by default but the BQF data structure does not impose any value.

#### 2.1.2 Hash function

A hash function is a mathematical transformation that takes an input (here a sequence of characters) and produces a number, called a hash value. In the current framework, the used hash function produces a value that is coded with a fixed-size number of bits. This transformation is designed to be deterministic, to produce an uniform distribution, and we want it to be as fast as possible. Given a hash function, two distinct elements are said in “collision” if they have the same hash value. A hash function is called “perfect” if it is injective, also meaning that there are no elements in collision. In this paper, we made the choice to use a xorshift [17] hash function, producing numbers between 0 and 2^2*k*^ for every *k*-mer. We use this [28] xorshift hash function as it is a Perfect Hash Function (PHF), preventing collisions. Also, as long as we project the *k*-mers into values of 2*k* bits, the function is reversible. It means that we can retrieve the original *k*-mer from its hashed value. The use of a non perfect hash function would also be possible but would imply the impossibility to enumerates the elements entered in the data structure. This would prevent, for example, the resizing of the data structure. As we want a fully dynamic data structure, we made the choice of using a PHF.

### 2.2 Quotient Filter (QF)

The work we present, called the Backpack Quotient Filter (BQF), is based on the Quotient Filter (QF) structure [3]. In this section, we provide a brief overview of the fundamental aspects of the QF structure, which is essential for comprehending our contribution. A deeper description of the structure is proposed in the Supplementary Materials.

A QF is a data structure that is used to store a set of elements. It is composed of a table with 2^*q*^ slots, each of fixed size *r*, where *q* and *r* are initially defined by the user. *q* and *r* are subject to change as the size of the table may change. It utilizes a hash function *h* that hashes elements to integers of *q* + *r* bits. When an element *x* is inserted, its hash value *h*(*x*) is computed and split into two parts:

- *h*_0_(*x*) of size *q* bits, called the “quotient”. It is used as an address in the table;
- *h*_1_(*x*) of size *r* bits, called the “remainder”. It is a fingerprint and is effectively stored in memory. *h*_1_(*x*) is inserted at the address *h*_0_(*x*).

To query the presence of an element *y* in the structure, *h*_0_(*y*) and *h*_1_(*y*) are computed. Finding *h*_1_(*y*) at position *h*_0_(*y*) induces that *y* is present. Figure 1 pictures the insertion step at slot 3, where solid hatched green lines symbolize the *r* bits of the remainder, inserted at address 3.

**Figure 1.**
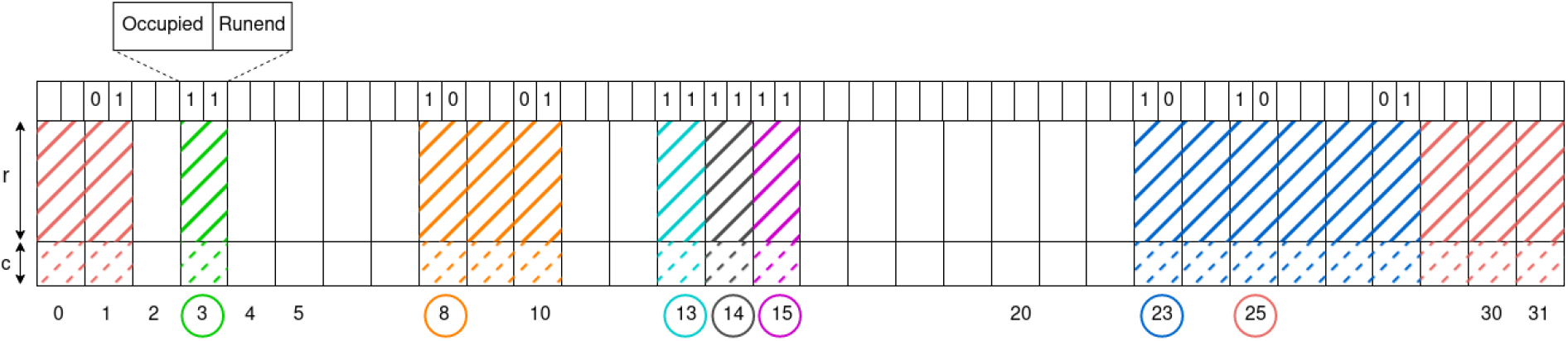
A 32 slots long BQF (*q*=5). First line represents the metadata bits (see [19] for more details). This short example does not represent blocks (*cf*. supplementary data) in the BQF for simplicity. Each slot has a size of *r* bits for the remainder with *c* bits for counts and 2 bits of metadata: *occupied* and *runend*. A circled address *Q* means that at least one element *x* such that *h*_0_(*x*) = *Q* has been inserted. Multiple remainders sharing the same color in the BQF have been originally inserted at the same address and form a run (*cf*. supplementary data). Empty metadata bits are set to 0.

Originally the QF is said to be a probabilistic data structure: with a non-null false-positive rate when querying elements. In the current framework, as we use a PHF, when querying a stored element, the false-positive rate is zero. However, as explained later (Section 2.3.2), we use an additional technique that does not exactly query the actual stored elements, and that generates a negligible but non-null false positive rate at query time.

In practice, the QF implementation we use is based on the Rank & Select Quotient Filter first proposed in [19]. We detail this operation in the Supplementary Materials.

#### 2.2.1 Abundance in Quotient Filters

As previously defined, the Quotient Filter structure is enough to handle the presence or absence of *k*-mers. It is possible to adapt the structure so it can store each *k*-mer alongside with its abundance in the indexed dataset. The Counting Quotient Filter [19] (CQF) is an example of a QF with abundance.

In the CQF, the abundance of each inserted element can be stored using the following process. A slot is used to store a remainder or an abundance value. If an element *x* has its abundance 1 ≤ *n* ≤ 2, then the element is inserted *n* times (with *n* = 2, two consecutive slots store *h*_1_(*x*)). When *n >* 2, *h*_1_(*x*) is stored twice, forming boundaries in the table, and *n* − 2 is encoded and stored between both boundaries, using potentially two slots or more. An extra slot holding a 0 might be necessary to maintain consistency in the runs. The point here is that this approach uses 2 slots when *n* = 2 and 3 or more when *n >* 2.

In the following section, we present our contribution, called the Backpack Quotient Filter, improving both the way the counts are stored and highly optimizing the size taken by each element to be queried.

### 2.3 The Backpack Quotient Filter

#### 2.3.1 Storing the abundance

In the BQF, the abundance of any element is stored using the following approach. As represented in Figure 1 each slot stores both a remainder and an abundance value. More precisely, each slot stores *r* bits for the remainder, and *c* extra bits are used to encode the abundance value. The *c* parameter is a user-defined parameter. The choice of *c* has a direct impact on the BQF size, adding *c×*2^*q*^ bits. The maximum value for abundance is 2^*c*^ and the value can be an exact count, or an order of magnitude (e.g. encoding of *log*_2_ values), offering flexibility based on precision requirements.

Compared to the CQF, the BQF uses only one slot per element, regardless of the abundance of this element, but at the same time it uses more bits per slot. However, thanks to the proposition described in the next section, we can reduce the size of the stored remainder. In this way, we cancel out this effect, even using less space per element while storing abundances.

#### 2.3.2 Reducing the space usage

In order to reduce the space usage, we take advantage of a method called “Fimpera” [26]. This method is originally designed to reduce the false-positives of data-structures having non-null false positive rates. Focussing on the presence/absence only, the key idea can be summarized as: if a word is present in a text, then all of its sub-words are present. Conversely, if any sub-word is absent, then the whole word is absent. In practice, instead of indexing the *k*-mers from a dataset, we insert all its *s*-mers, with *s* ≤ *k*. At query time a *k*-mer is considered as indexed if and only if all its *s*-mers are indexed in the structure. In the general case of querying a *k*-mer in an structure with hard collisions, this approach enables to lower the false positive rate of the query because all *s*-mers of a specific *k*-mer need to be false positives to create a false positive *k*-mer.

The same idea can be exploited when taking the abundance into account. The abundance of a *k*-mer is at most equal to the least abundant *s*-mer it is composed of. Therefore, we store the abundance of s-mers in the filter and report the abundance of a queried k-mer as the minimum of the abundances of the s-mers composing it. The techniques described in [26] explain how this approach does not have a negative impact on query time and may even improve it. When applied to a structure having hard collisions, this approach limits the overestimation of the abundance, as all the *s*-mers of a queried *k*-mer have to be overestimated to overestimate the real abundance of this *k*-mer.

In the BQF, we do not have any hard collision. We apply this approach to gain space instead.

Let us first study the size of the reversible hash value, used to store words on a four-character alphabet. Each character (here {*A, C, G, T* }) requires two bits for its encoding. Hence, encoding a word of length *l* requires 2*l* bits. As we use a reversible hash function, the size of the hash value requires the same size as the original encoded data, 2*l*.

By inserting *s*-mers, smaller than *k*-mers, the size of the reversible hash value of each inserted element becomes 2*s* instead of 2*k*. If we denote by *z* the difference between *k* and *s*, the gain is 2*z* bits per element. In the BQF structure, the consequence is that the size of each slot is decreased by 2*z*. All in all, applying this approach enables to save 2^*q*^ *×* 2*z* bits. The same PHF is used, with the same properties of injectivity and reversibility of stored elements.

A drawback of using this approach is the lost of the enumerating feature for *k*-mers. The hash function is still reversible but because we have *s*-mers in the filter, we can only reconstruct (and thus enumerate) these *s*-mers and not the *k*-mers we want to query. It is important to note that we only lose the *k*-mers enumeration, not the dynamicity: resizing the BQF remains possible.

The second drawback of applying this approach is the creation of a new kind of false positives, called “construction false positives”. The existence of construction false positive is explained by a simple sentence: a *k*-mer may be absent but all of its *s*-mers may be present. We meet this case if each *s*-mer of an absent *k*-mer *x* has been individually inserted through the present *k*-mers sharing *s*-mers with *x*.

#### 2.3.3 Theoretical influence of the *s* parameter

We now detail the theoretical consequences of reducing the size *s* of indexed elements, with *s* ∈]0, *k*].

1. Decreasing *s* increases the “construction false positive” rate. The smaller the *s* value is, the higher is the probability that a queried *k*-mer, non existing in the indexed set, has all its *s*-mers existing in this set.
2. Decreasing *s* may increase the number of indexed *s*-mers in short reads datasets. A sequence of size *l* contains (*l* − *k* + 1) *k*-mers and (*l* − *s* + 1) *s*-mers. Hence, it contains *z* = *k* − *s* additional *s*-mers than *k*-mers. This is negligible while indexing for instance an assembled genome. But when it comes to index millions of reads with low redundancy between them, as this is the case in our experimentations using sea-water metagenomes, each of the million reads contains *z* more *s*-mers than *k*-mers, with a low redundancy between reads.
3. Decreasing *s* decreases the size taken by each indexed *s*-mer, which is the expected effect. This is the main advantage of the approach. Recall that the total size of structure is reduced by 2^*q*^*×* 2*z* when using *s*-mers instead of *k*-mers. Hence the smaller *s* is, the more space is saved.

In general, the results presented Section 3.3 suggest that the size of the data structure decreases as *s* decreases, despite the conflicting effects of the last two previous points. Selecting small *s* values only has the potential to increase the construction false positive rate. However, when using recommended values, it stays below 10^*−*5^%.

#### 2.3.4 Doubling the number of slots when the structure is full

One of the main advantage of building the QF with a PHF is that conversely to a Bloom filter for instance, when the structure is full, it is possible to double its number of slots (from *q* to *q* + 1). During this process, the hash value of each element remains the same, but the way it is distributed between the quotient and remainder changes. This occurs because, after doubling, *q* + 1 bits are used to represent the address, while *r* − 1 bits are used for the remainder. Finally, the total number of stored elements faces no theoretical limitation.

In practice, for performances reasons, one doubles the number of slots when the “load factor” (number of stored elements divided by the number of slots) becomes bigger than 95%.

#### 2.3.5 Number of bits per stored element

As stated, the basis of the QF data structure is to use the address of stored elements as a part of their hash value. As a consequence, the size of the remainder stored for each element decreases when the number of slots increases. This is not linear. Let us consider the initial scenario, where the QF is composed of 2^*q*^ slots in which *r* bits per slot are used as remainder. In this case, the BQF uses 2^*q*^ *×*(*r* + *c* + 3) bits, as for each stored element, *r* bits store the remainder, *c* bits are used to store the abundance, and 3 additional metadata bits are used by the structure itself (*runend, occupied* and a third one, *offset*, explained in supplementary data). In this situation, if this structure is full, each of the 2^*q*^ stored elements requires (*r* + *c* + 3) bits.

Now consider that the size of the structure doubles in order to index more elements. The structure then contains 2^*q*+1^ slots. In this situation, *q* + 1 bits indicate the address of each slot, and so the remainder of each element decreases to *r* − 1 bits instead of *r*. In this case, the total size of the structure becomes 2^*q*+1^ *×* (*r* − 1 + *c* + 3) = 2^*q*+1^ *×* (*r* + *c* + 2). When this structure is full, each of the 2^*q*+1^ stored elements now requires (*r* + *c* + 2) bits instead of (*r* + *c* + 3). By doubling again the size of the structure, it would contains 2^*q*+2^ slots, each composed of (*r* + *c* + 1) bits. When this structure in turn becomes full, each element require (*r* + *c* + 1) bits, and so on.

Note that this practical effect ends when the remainder is empty, in which case the full hash value of each element is entirely given by the address of the element. This is a theoretical view as, in the case of *k*-mer indexing, when indexing conventional *k* value typically around 30, approximately 140 petabytes would be needed to contain the 4^*k*^ slots (the number of possible distinct *k*-mers).

## 3 Results

We propose some experiments on real metagenomic datasets. The objective is to compare the performances obtained with the BQF with those obtained using state-of-the-art data structures for indexing *k*-mers together with their abundances, based on the Quotient Filter: the CQF [19] and on the counting Bloom Filter: the CBF [26]. We also included a comparison with Bifrost [12] and SSHash [22]. Both of these approaches allow for querying indexed *k*-mers, but they have significant differences in their main features, which are summarized below.

These results also enable to show the impact of the unique parameter introduced by BQF: the *s* value. We also show the influence of the number of indexed elements on the whole data structure size.

The version used for BQF is v1.0.0. Details about protocols and links to datasets are available online [1].

### 3.1 Used datasets

Our results were obtained on three distinct metagenomic datasets in which we exclusively considered *k*-mers present two or more times.

- Dataset “*sea-water34M* ”: 34 million Illumina reads from the *Tara* Oceans sequencing project. The compressed *fastq* file is 7.7GB. It contains 257M distinct 32-mers and 346M distinct 19-mers occurring at least twice.
- Dataset “*sea-water143M* ”: 143 million Illumina reads from the *Tara* Oceans sequencing project. The compressed *fastq* file is 33GB. It contains 1.2B distinct 32-mers and 1.5B distinct 19-mers occurring at least twice.
- Dataset “*gut* ”: 13 million reads from pig microbiota Pacbio sequencing. The compressed *fastq* file is 42GB. It contains 475M distinct 32-mers and 420M distinct 19-mers occurring at least twice.

These sea-water and gut microbiota metagenomic datasets are representative of a highly complex situation, with a large diversity content. For instance, there are 9.5 billion *k*-mers in *sea-water143M* dataset, leading to a set of 5.7 billion distinct *k*-mers. Among them, only 1.2 billion are present twice or more. For the *gut* dataset, we counted 22 billion *k*-mers, 1.2 billion distinct ones and 0.475 billion are present twice or more. We propose in Supplementary Materials a visualisation of the *k*-mer spectrum of the sets *sea-water143M* and *gut*. They illustrate the complexity of these datasets, where there is no peak linked to the specific presence of one or more species.

### 3.2 Compared performances

In this section, we propose to compare the BQF with the CQF (https://github.com/splatlab/cqf, commit 68939f5). Both structures use the same Xorshift hash function, a PHF, ensuring no collisions. We also compare with results obtained with a counting Bloom filter (CBF) implementing the Fimpera approach (https://github.com/lrobidou/fimpera, commit 662328d). Both CBF and BQF use 5 bits for counters (*c* = 5), allowing a maximal abundance value of 64 as we store exact values. BQF and CBF use the Fimpera approach, initialized with *k* = 32 and *s* = 19, thus 19-mers are counted and inserted. The sizes of the BQF and CQF are determined solely by the total number of elements plus the element abundances for the CQF. Regarding the CBF, we decided to create a CBF of the same size as the BQF. This ensures fair comparisons when considering a fixed amount of disk space. The choice of parameters is discussed further in this section.

We also show results obtained by Bifrost (version 1.3.1) and SSHash (version 3.0.0). Those comparisons are not exactly fair as these tools embed additional features (computing pre-assembly of the data in the so-called compacted de Bruijn graph, possibly indexing multiple datasets for Bifrost) while Bifrost cannot index the abundance, and while SSHash is a static data structure. However, it is interesting to present these results as they show that these state-of-the-art tools —which are not specifically designed for the task of only indexing *k*-mers with their abundances— are not optimal for this task.

#### 3.2.1 Experimental setup

We computed the build time and the query time for each approach. In addition to the building time, the results show the pre-processing time, *i.e*. the time used to obtain the correct input file from the raw compressed *fastq* file (counted *k*-mers for CQF, CBF and BQF, and SPSSs [24] for SSHash).

The parameters are *k* = 32 for CBF, CQF, and BQF, *c* = 5 (counters size for BQF, CBF), and *s* = 19 (19-mers were inserted for BQF, CBF). For SSHash and Bifrost, we used *k* = 31 as using *k* = 32 would have doubled the *k*-mer encoding size for Bifrost, and because SSHash uses *k* ≤ 31. Bifrost used 4 threads and *m* = 17 for SSHash (minimisers size).

Positive queries in a dataset 𝒟 are *k*-mers reads from 𝒟 itself. Negative queries are *k*-mers from randomly generated sequences (between 80 and 120 nucleotides). Around 2 billion *k*-mers over 30 million sequences were positively queried. Around 7 billion negative *k*-mers over 100 million sequences were negatively queried.

BQF and CQF sizes are measured experimentally. Their size corresponds exactly to their theoretical value, also showing that, thanks to the simplicity of the structure, no space overhead is required. CBF size was chosen to be the same as BQF’s. SSHash size is the one displayed by the tool at the end of the building step. Bifrost size is measured as the peak memory usage after loading the graph and the index in memory (from binary representation on disk).

The executions were performed on the GenOuest platform on a node with 4 *×* 8 cores Xeon E5-2660 2,20 GHz with 128 GB of memory.

#### 3.2.2 Comparative results

##### Comparing CQF and BQF

Compared to the CQF, the major advantage of the BQF is in term of space. As shown Table 1, the BQF is approximately four times smaller than the CQF for every indexed dataset. The same advantage is found in terms of space efficiency (bits/element), being approximately 5 to 7 times more efficient. However, one drawback is the occurrence of false positive calls, which are generally less than 10^*−*5^% and can even be as low as 0% in the gut data set.

**Table 1:** Comparative performances. Recall that Bifrost and SSHash do not index the same number of elements than CQF, CBF and BQF, explaining the difference in term of number of bits per element as compared to the structure size. Given its computation time (≥ 24 hours on the *sea-water143M* dataset), we report SSHash results only for the *sea-water34M* dataset.

| Dataset | Structure | Memory (GB) | bits per element | Pre-processing + Build time (s) | Pos. query time (kmer/s) | Neg. query time (kmer/s) | FP rate (%) |
| --- | --- | --- | --- | --- | --- | --- | --- |
| *sea-water34M* | Bifrost | 3.67 | 111 | 871<sup>α</sup> | 3,823,877 | 1,767,818 | 0 |
|  | SShash | 0.39 | 12 | 1,626<sup>β</sup> + 34,890 | 1,460,401 | 1,450,285 | 0 |
|  | CQF | 4.85 | 151 | 215<sup>γ</sup> + 210 | 3,267,592 | 4,110,300 | 0 |
| | CBF | 1.14 | 26 | 219<sup>γ</sup> + 429 | 55,852 | 62,229 | $4.8 \times 10^{-6}$ |
| | **BQF** | 1.14 | 26 | 219<sup>γ</sup> + 257 | 3,090,526 | 3,506,136 | $1.6 \times 10^{-6}$ |
| *sea-water143M* | Bifrost | 17.59 | 115 | 4,252<sup>α</sup> | 1,977,587 | 1,766,008 | 0 |
|  | CQF | 9.70 | 129 | 771<sup>γ</sup> + 949 | 3,254,682 | 3,931,239 | 0 |
| | CBF | 4.03 | 24 | 780<sup>γ</sup> + 2,039 | 55,044 | 61,407 | $5.8 \times 10^{-5}$ |
| | **BQF** | 4.03 | 24 | 780<sup>γ</sup> + 1,101 | 2,601,710 | 2,562,646 | $3.0 \times 10^{-5}$ |
| *gut* | Bifrost | 5.85 | 99 | 6,137<sup>α</sup> | 4,003,868 | 1,653,834 | 0 |
|  | CQF | 19.39 | 158 | 1,178<sup>γ</sup> + 396 | 2,527,472 | 3,666,312 | 0 |
| | CBF | 1.14 | 22 | 1,085<sup>γ</sup> + 468 | 56,009 | 61,356 | $1.6 \times 10^{-6}$ |
|  | **BQF** | 1.14 | 22 | 1,085<sup>γ</sup> + 341 | 2,855,172 | 3,317,307 | 0 |

<sup>α</sup> Bifrost does not require pre-processing step
<sup>β</sup> BCALM [8] (https://github.com/GATB/bcalm, version 2.2.3) for unitigs + UST [24] (https://github.com/jermp/UST, commit b3d0710) for simplitigs
<sup>γ</sup> KMC [13] (https://github.com/refresh-bio/KMC, version 3.2.4) kmer counting

##### Comparing CBF and BQF

The results presented Table 1 indicate that the false-positive rate is slightly better with the BQF compared to CBF. However, both approaches still have a very low false positive rate of approximately 10^*−*5^%, which is insignificant for indexing and pseudo-alignment applications. BQF offers several significant benefits over CBF. First, BQF allows for faster time queries, with an average speed improvement of 50 times compared to CBF. Additionally, BQF does not have any theoretical limitations on the number of stored elements, unlike CBF which is designed for a fixed maximum number of elements that cannot be updated. Finally, the elements stored in a BQF (the *s*-mers) can be enumerated, while this is not the case with the CBF.

##### Abundance overestimation due to the Fimpera approach

In this work, we did not recompute the so-called overestimation inherent to the Fimpera abundance representation. This overestimation is in the order of 1% to 2% according to the results presented in [26], meaning that 1 to 2% of the abundances of true positive calls are overestimated. Furthermore, for those results that were overestimated, the average difference was shown to be approximately 1.07 times the correct abundance range. All in all, this slight overestimation, limited to less than 2% of the calls, has no significant impact while estimating the abundance of a query composed of at least dozens or hundreds of *k*-mers.

##### Bifrost and SSHash results

As shown by results presented Table 1, Bifrost is approximately two times slower than BQF to build the data structure and more than twice as slow to perform negative queries. It uses approximately 4.5 times more space per element, and more importantly, it does not provide the abundance of *k*-mers. The SSHash approach, for its part, taking advantage of super-*k*-mers [15], uses approximately 2 times less space per element than BQF. However, it is static and is nearly two orders of magnitude slower to construct, drastically limiting its application to large-scale projects.

### 3.3 Impact of size of the indexed *s*-mers

As stated earlier, the BQF structure stores *s*-mers, to emulate *k*-mers at query time, with *s* ≤ *k*. The choice of the *s* value has several consequences that are described Section 2.3.3, and that we propose to empirically observe here.

#### 3.3.1 Effect of *s* on the number of construction false positives

Figure 2 illustrates that using any value of *s* bigger than 17 enables to limit drastically the construction false positive rate. When *s*-mers become smaller than 17 nucleotides, the probability they appear by chance in sequences composed of millions of characters on the { *A, C, G, T* } alphabet becomes close to 1. In this case, mostly any *k*-mers can be constructed from these *s*-mers, explaining the nearly 100% construction false positive rate.

**Figure 2.**
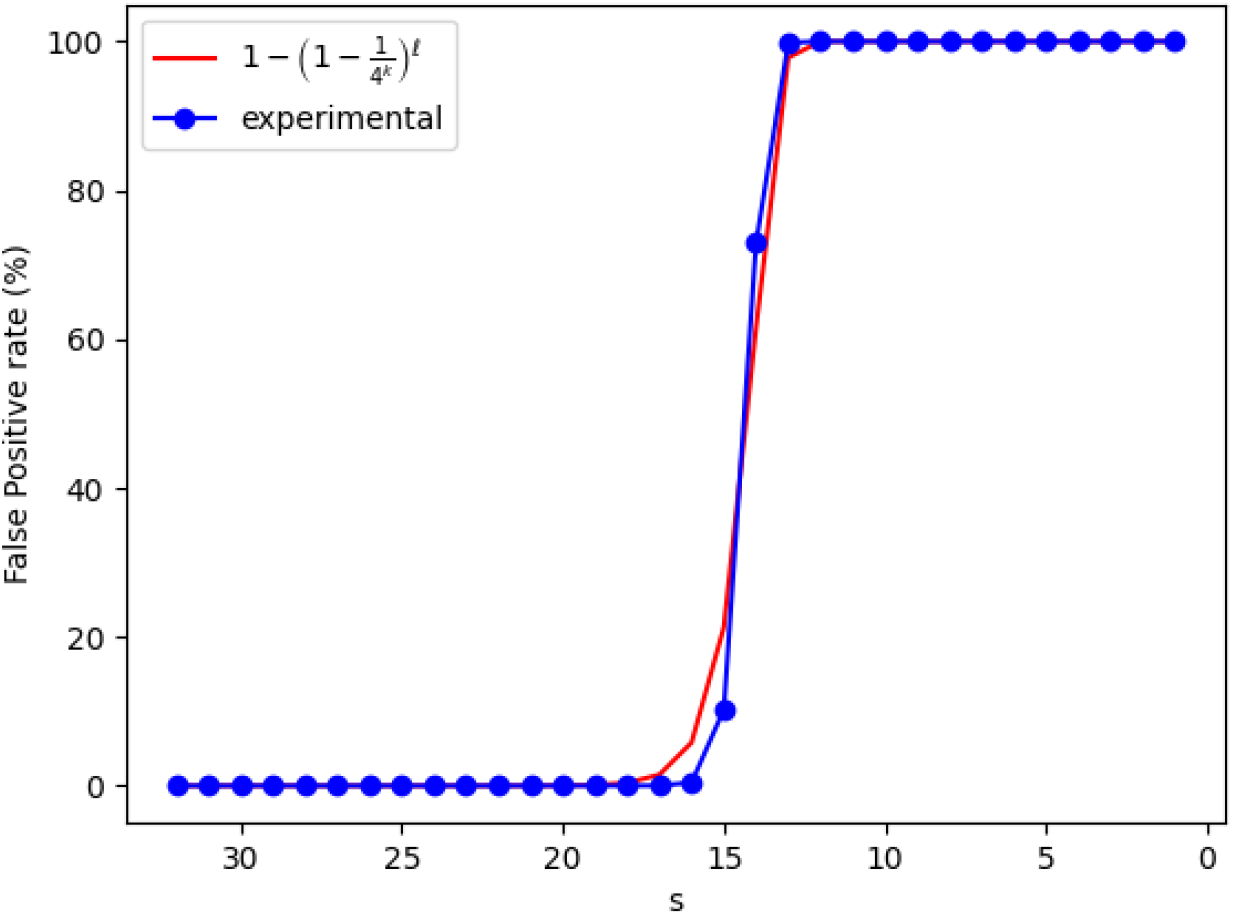
Empirical observation of the evolution of the construction false positive rate with respect to *s*. Indexed dataset: “*sea-water34M* ”, querying random 32-mers

Note that the shape of this curve is highly correlated with the probability that an element of size *s* appears by chance in a sequence of size *l*. This probability is equal to 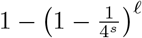. In concrete terms, this allows a user to reliably determine a value of *s* knowing *l*, even approximately. The value of *l* can be approximated thanks to the number of distinct *k*-mers in the dataset (as this is the case in Figure 2), efficiently computed by ntCard [18] for instance.

#### 3.3.2 Effect of *s* on the structure size

Recall that decreasing *s* has two opposite effects on the structure size:

(a) in certain conditions (see below), decreasing *s* can increase the number of indexed *s*-mers, which tends to increase the size of the structure (need to double its size when reaching 95% load factor);
(b) decreasing *s* decreases the remainder size, and so decreases the total size of the structure.

In this section, we propose to observe the practical consequences of this choice.

(a) Figure 3 shows (plain blue curve) the number of distinct *s*-mers according to *s*. With long enough *s*-mers (*s >* 17), decreasing *s* sub-linearly increases the number of distinct *s*-mers. This is true in the case of relatively short reads, with new generation sequencing for instance (Illumina example within Figure 3 with *sea-water34M* dataset). On the other hand, third-generation sequencing produces longer reads, in this context decreasing *s* decreases the number of elements to index (475M distinct 32-mers and 420M distinct 19-mers in *gut* PacBio dataset). Table 1 shows this result: when comparing BQF and CQF building time (which depends on the number of elements to index), we can see that BQF is slightly faster on *gut* (PacBio) dataset as there are fewer 19-mers than 32-mers.

**Figure 3.**
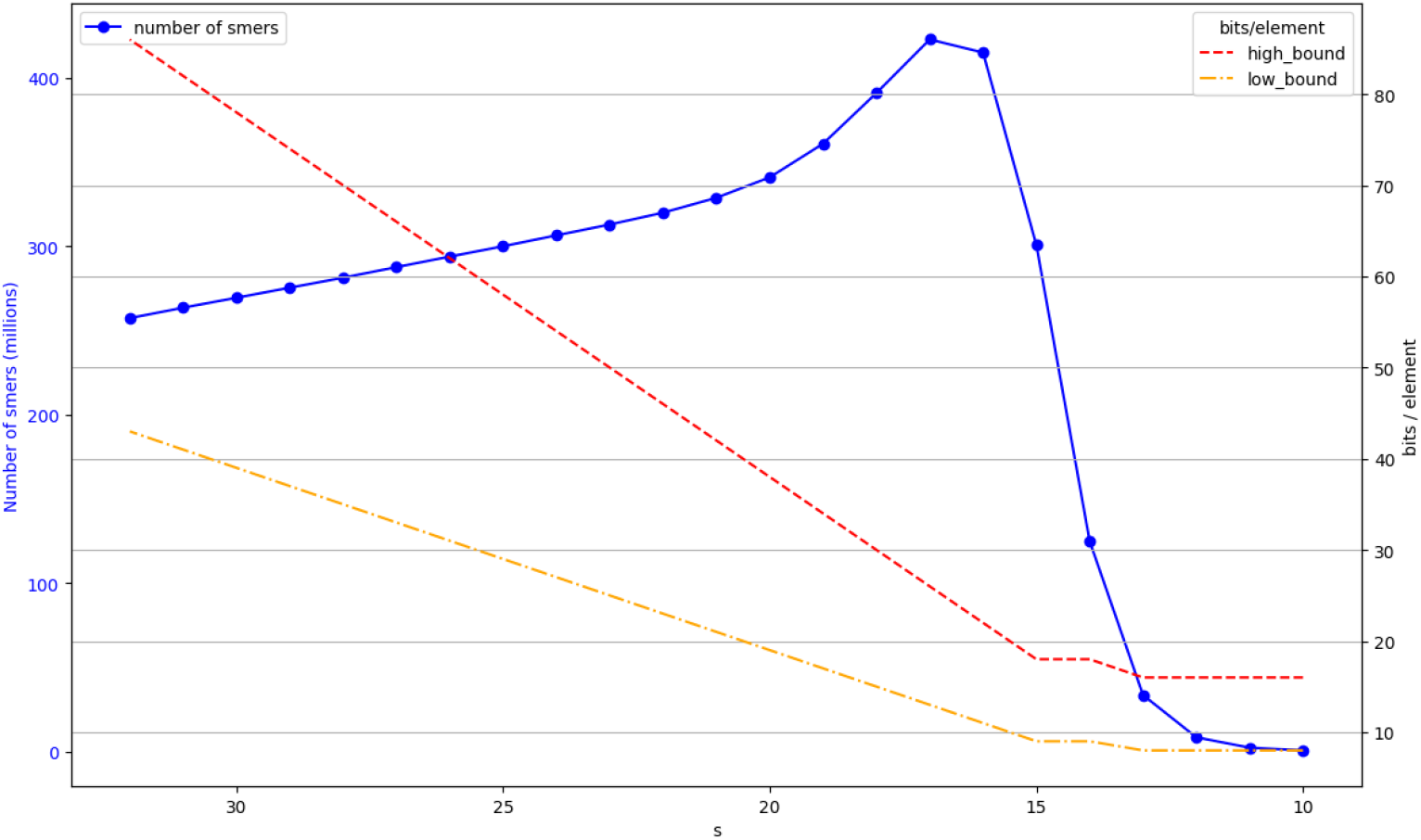
Evolution of the number of *s*-mers depending on *s* in an Illumina sequencing dataset (*sea-water34M*): plain blue line. Evolution of the number of bits per element depending on *s* on the same dataset. *high bound* is the red upper dotted curve, corresponding to a half-full BQF. *low bound* is plotted in orange, under *high bound* and corresponds to a full BQF (95% load factor)

With *s* ≤ 17, another effect exists: nearly all the *s*-mers exist in the text, and so the number of distinct *s*-mers becomes limited by 4^*s*^, explaining why the number of distinct *s*-mers decreases when *s* decreases below *s* = 17.

(b) The two dashed lines of Figure 3 show the number of bits per element either if the structure is half full or considered as full (in practice one doubles the structure size if its load factor is 95%). The observation is that even on this highly complex sea-water metagenomic dataset, the space needed to store *s*-mers decreases when *s* decreases, even though more *s*-mers have to be stored. A fictitious example is available in the supplementary data, Section “Side effect of lowering *s*”, demonstrating a case where the number of *s*-mers reaches a doubling threshold before the number of *k*-mers. Results show that, even in this case, the high and low bound of number of bits per element, is never increasing while *s* decreases.

All in all, regarding the data-structure size, the best choice is to use *s* as small as possible, but bigger than 17 to avoid an explosion of the construction false positive rate, as it keeps it below 10^*−*5^% in this setup.

### 3.4 Effect of the number of indexed elements on the structure size

Based on our metagenomic samples, this section comments on the experimental value of bits per element (see section 2.3.5) used by the BQF compared to CQF. Figure 4 shows the evolution of the data structure size (A) and the evolution of bits per element (B) while elements are inserted.

**Figure 4.**
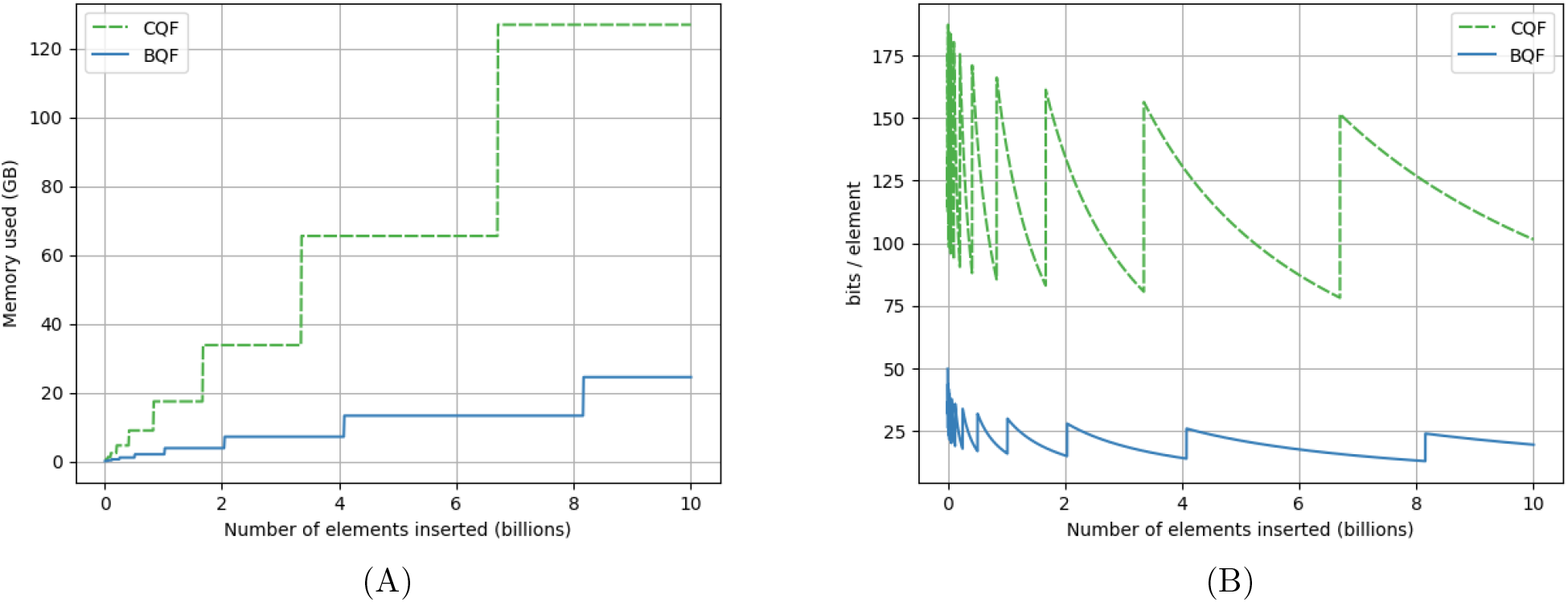
Effect of the number of indexed elements on the size and space efficiency. Generated from indexing dataset “*sea-water34M* ”, *k* = 32, *s* = 19 and *c* = 5 for BQF. **(A)** total data-structure size. **(B)** size in terms of number of bits per indexed element.

The stairs shapes of figure 4(A) are due to the size of the data structure that doubles each time their load factor reaches 95%. Then, each insertion increases the load factor without consuming more space. The figure highlights the fact that on real metagenomic datasets, the CQF needs a lot of space due to the counter encoding which uses an average of 2.44 slots per element (in the *sea-water34M* dataset). Given a fixed number of insertions, because the CQF doubles its size 2.44 times more frequently, the total size occupied by the CQF is much higher than that of the BQF. At least with metagenomics data, while counts are low but not unique, BQF will always occupy less space than CQF.

In figure 4(B) the dented curves show the space used per element. The curves are decreasing as the data structures are filled with elements. The vertical jumps correspond to the data structure resizes. We can see that the two structures behave in the same way while the BQF uses fewer bits per element. It is explained by the number of slots per element (a 2.44 times decrease) but also by the Fimpera scheme used in the BQF approach. An interesting fact is that the peaks for both structures get lower while the data structure size doubles. This is because the slots are one bit shorter after each resize, as explained Section 2.3.5.

Finally, at the price of a negligible non-null false positive rate (in the order of 10^*−*5^% to 10^*−*6^% in our experiments), the BQF enables to make queries among dozen of billions of elements, using between 13 and 26 bits per element, while the CQF requires between 75 and 150 bits per element for the same settings.

## 4 Conclusion

This paper introduces the Backpack Quotient Filter, a quotient filter with abundance. The BQF, like other quotient filters, uses a table to store elements. Only a fraction of these elements is explicitly stored, as the rest is implicitly given through their address. Specifically in the BQF, for every element, *c* additional bits are used to encode the abundance associated. This strategy enables to index billions of elements with their abundance using between 13 and 26 bits per element, depending on the data structure load factor.

In addition to this counting strategy, the BQF implements the Fimpera strategy, which emulates *k*-mers from their *s*-mers (with *s* ≤ *k*). A direct consequence of this emulation is a gain of 2^*q*^*×*2*×*(*k* − *s*) bits over the whole structure, with 2^*q*^ being the number of slots in the BQF. Our results show that the results are robust with respect to the *s* parameter, as long as *s* is bigger than a fixed threshold, namely *s >* 17.

Our results indexing metagenomic data show that the BQF size is at least four times more compact than the most similar data structure: the Counting Quotient Filter [19]. The indexing and query times are in the same order of magnitude. This result is at the price of a non-null but extremely low false positive rate (≈ 10^*−*6^% in our experiment). To fully benefit from the flexible sizes of the counters, if the user can afford it, it is advised to index orders of magnitude (e.g. *log*_2_ values) instead of exact counts.

The BQF inserts hash values of the elements. By using a perfect hash function, we ensure having no collisions among stored elements. This offers the possibility to enumerate the elements stored in the structure. If the structure gets full when adding elements, this offers a way to relocate all elements after doubling the size of the structure. So there is no theoretical limit to the number of elements stored in the BQF. This dynamicity is significant in the context of intensive sequencing and indexing.

## Funding

The work was funded by the Inria Challenge “OmicFinder”, the ANR SeqDigger (ANR-19-CE45-0008), and the European Union’s Horizon 2020 research and innovation program under the Marie Sklodowska-Curie grant agreement No 956229. The funders had no role in study design, data analysis, decision to publish, or manuscript preparation.

## Acknowledgements

We acknowledge the GenOuest bioinformatics core facility https://www.genouest.org for providing the computing infrastructure. The authors thank EÉmeline Roux for providing the link to the *gut* dataset.

## Notes

https://github.com/vicLeva/bqf

### Competing Interest Statement

The authors have declared no competing interest.

## References

[1] Experiments details and protocols. https://github.com/vicLeva/bqf/wiki/ Experiments-details-and-protocol-for-BQF-paper-results.

[2] Jarno N Alanko, Jaakko Vuohtoniemi, Tommi Mäaklin, and Simon J Puglisi. Themisto: a scalable colored k-mer index for sensitive pseudoalignment against hundreds of thousands of bacterial genomes. Bioinformatics, 39(Supplement 1):i260–i269, June 2023.

[3] Michael A. Bender, Martin Farach-Colton, Rob Johnson, Russell Kraner, Bradley C. Kuszmaul, Dzejla Medjedovic, Pablo Montes, Pradeep Shetty, Richard P. Spillane, and Erez Zadok. Don’t Thrash: How to Cache Your Hash on Flash, August 2012. arXiv:1208.0290 [cs].

[4] Timo Bingmann, Phelim Bradley, Florian Gauger, and Zamin Iqbal. COBS: A Compact Bit-Sliced Signature Index. In Nieves R. Brisaboa and Simon J. Puglisi, editors, String Processing and Information Retrieval, Lecture Notes in Computer Science, pages 285–303, Cham, 2019. Springer International Publishing.

[5] Burton H. Bloom. Space/time trade-offs in hash coding with allowable errors. Communications of the ACM, 13(7):422–426, 1970.

[6] Phelim Bradley, Henk C. den Bakker, Eduardo P. C. Rocha, Gil McVean, and Zamin Iqbal. Ultrafast search of all deposited bacterial and viral genomic data. Nature Biotechnology, 37(2):152–159, February 2019. Number: 2 Publisher: Nature Publishing Group.

[7] Josephine Burgin, Alisha Ahamed, Carla Cummins, Rajkumar Devraj, Khadim Gueye, Dipayan Gupta, Vikas Gupta, Muhammad Haseeb, Maira Ihsan, Eugene Ivanov, et al. The european nucleotide archive in 2022. Nucleic Acids Research, 51(D1):D121–D125, 2023.

[8] Rayan Chikhi, Antoine Limasset, and Paul Medvedev. Compacting de Bruijn graphs from sequencing data quickly and in low memory. Bioinformatics (Oxford, England), 32(12):i201–i208, June 2016.

[9] Andrea Cracco and Alexandru I. Tomescu. Extremely fast construction and querying of compacted and colored de Bruijn graphs with GGCAT. Genome Research, 33(7):1198–1207, July 2023.

[10] Thomas Mueller Graf and Daniel Lemire. Xor Filters: Faster and Smaller Than Bloom and Cuckoo Filters. ACM Journal of Experimental Algorithmics, 25:1–16, December 2020. arXiv:1912.08258 [cs].

[11] Thomas Mueller Graf and Daniel Lemire. Binary Fuse Filters: Fast and Smaller Than Xor Filters. ACM Journal of Experimental Algorithmics, 27:1–15, December 2022. arXiv:2201.01174 [cs].

[12] Guillaume Holley and Páall Melsted. Bifrost: highly parallel construction and indexing of colored and compacted de Bruijn graphs. Genome Biology, 21(1):249, September 2020.

[13] Marek Kokot, Maciej Dlugosz, and Sebastian Deorowicz. KMC 3: counting and manipulating k-mer statistics. Bioinformatics, 33(17):2759–2761, September 2017.

[14] Téeo Lemane, Paul Medvedev, Rayan Chikhi, and Pierre Peterlongo. kmtricks: efficient and flexible construction of Bloom filters for large sequencing data collections. Bioinformatics Advances, 2(1):vbac029, January 2022.

[15] Yang Li, Pegah Kamousi, Fangqiu Han, Shengqi Yang, Xifeng Yan, and Subhash Suri. Memory efficient minimum substring partitioning. Proceedings of the VLDB Endowment, 6(3):169–180, January 2013.

[16] Camille Marchet and Antoine Limasset. Scalable sequence database search using partitioned aggregated Bloom comb trees. Bioinformatics, 39(Supplement 1):i252–i259, June 2023.

[17] George Marsaglia. Xorshift RNGs. Journal of Statistical Software, 8:1–6, July 2003.

[18] Hamid Mohamadi, Hamza Khan, and Inançc Birol. ntcard: a streaming algorithm for cardinality estimation in genomics data. Bioinformatics, 33:1324–1330, 2017.

[19] Prashant Pandey, Michael A. Bender, Rob Johnson, and Rob Patro. A General-Purpose Counting Filter: Making Every Bit Count. In Proceedings of the 2017 ACM International Conference on Management of Data, SIGMOD ’17, pages 775–787, New York, NY, USA, 2017. Association for Computing Machinery.

[20] Prashant Pandey, Michael A Bender, Rob Johnson, and Rob Patro. Squeakr: an exact and approximate k-mer counting system. Bioinformatics, 34(4):568–575, February 2018.

[21] Giulio Ermanno Pibiri. Sparse and skew hashing of K-mers. Bioinformatics, 38(Supplement 1):i185– i194, June 2022.

[22] Giulio Ermanno Pibiri. On weighted k-mer dictionaries. Algorithms for Molecular Biology, 18(1):3, June 2023.

[23] Giulio Ermanno Pibiri and Roberto Trani. PTHash: Revisiting FCH Minimal Perfect Hashing. In Proceedings of the 44th International ACM SIGIR Conference on Research and Development in Information Retrieval, SIGIR ’21, pages 1339–1348, New York, NY, USA, July 2021. Association for Computing Machinery.

[24] Amatur Rahman and Paul Medvedev. Representation of k-mer sets using spectrum-preserving string sets. In Research in Computational Molecular Biology - 24th Annual International Conference, RECOMB 2020, Padua, Italy, May 10-13, 2020, Proceedings, volume 12074 of Lecture Notes in Computer Science, pages 152–168. Springer, 2020.

[25] Michael Roberts, Wayne Hayes, Brian R. Hunt, Stephen M. Mount, and James A. Yorke. Reducing storage requirements for biological sequence comparison. Bioinformatics (Oxford, England), 20(18):3363–3369, December 2004.

[26] Lucas Robidou and Pierre Peterlongo. fimpera: drastic improvement of Approximate Membership Query data-structures with counts, April 2023. Pages: 2022.06.27.497694 Section: New Results.

[27] Sanjay K. Srikakulam, Sebastian Keller, Fawaz Dabbaghie, Robert Bals, and Olga V. Kalinina. MetaProFi: an ultrafast chunked Bloom filter for storing and querying protein and nucleotide sequence data for accurate identification of functionally relevant genetic variants. Bioinformatics (Oxford, England), 39(3):btad101, March 2023.

[28] Thomas Wang. Integer hash function. https://gist.github.com/lh3/59882d6b96166dfc3d8d, January 1997.

