## Supplementary materials for "The Backpack Quotient Filter: a dynamic and space-efficient data structure for querying *k*-mers with abundance"

### The Backpack Quotient Filter: Supplementary Materials

#### Additional details about the CQF rank and select scheme and optimizations

Here we explain some key parts of the CQF, as originally proposed by Pandey *et.al.* in [1].

##### Soft Collision Resolution and Run Management

We differentiate two kinds of hash collision. One is called "hard collision", and happens when two distinct elements have the same hash value. We avoid this by using a PHF. The second is called "soft collision". It is inherent to quotient filters. A soft collision occurs when two distinct elements $x$ and $y$ have different hashes but the same quotient: $h_0(x) = h_0(y)$.

Because only one remainder can be inserted in any slot, additional remainders sharing the same quotient value are shifted into the next slots. Elements in soft collision are stored consecutively in the table, thus forming a so-called "run". Inside a run, the remainders are stored in ascending order. Formally, for all elements $x, y$ in a run, with $h_0(x) = h_0(y)$, and assuming that $h_1(x) < h_1(y)$, the slot address where $x$ is stored is lower than the one of $y$.

The slot address of the first element $x_{first}$ of a run may be distinct from $h_0(x_{first})$. The run can be shifted further than its insertion slot. This case appears when another upstream run already occupies the slot given by $h_0(x_{first})$.

Note that an element shifted from the last slot $(2^q - 1)$ goes into slot 0, since the structure is circular.

To keep track of the shifting process for later insertions and queries, two additional bits of metadata are used in each slot. They enable, thanks to rank and select operations, to determine the actual slot where an element is stored. Finally, an additional bit per slot is used to improve the theoretical complexity of the insertion and query and the practical speed.

- The *occupied* bit determines whether a slot welcomed an element, whose remainder could be shifted and located elsewhere. In Figure 1 (main text, Section Material & methods), slot 3 has its *occupied* bit set to one because the green run (1 element) has been inserted here. In slot 25, the occupied bit is also set to 1 because it is the insertion slot of the red run, even though the red run has been shifted further by the blue run.

- The *runend* bit indicates whether a slot stores a remainder that is at the end of a run. In this case, it is set to 1. Otherwise, it is set to zero, and in this situation, the next slot is either empty or is the first one of a different run.

In Figure 1 (Section Material & methods), 3 remainders are forming a run on slots 8, 9, 10. They all have been inserted with the quotient being 8 and then the biggest ones were shifted to the right. This simple run of 3 elements differs from slots 13, 14, 15 where we have 3 runs of 1 element each.

If we consider a CQF composed of $2^q$ slots, then we have two binary vectors: *occupieds* and *runends*, both of size $2^q$ bits. To find a possibly shifted run from a slot $i$, we aim to find the end of this run. One way to do so is by:

1. counting the number $d$ of runs that are present before $i$ by counting the number of 1 in *occupieds* before $occupieds[i]$. We call this operation $Rank(occupieds, i)$. $Rank(v, i)$ is defined as the number of 1's from position 0 to position $i$ (included) in a binary vector $v$.

2. finding the position of the $d^{th}$1 in *runends*. Here we have the second operation: $Select(runends, d)$, and we define $Select(v, i)$ = position of the $i^{th}$ 1 in the vector $v$.

All in all $runend\_position(i) = Select(runends, Rank(occupieds, i))$.

##### Block-based Optimization

When inserting or querying an element in the filter, the position of the run where it belongs needs to be computed. This means applying *Rank* and *Select* over *occupieds* and *runends*. As it becomes too expensive to iterate over a billion bits long vectors, the structure is divided into blocks of 64 slots. Each block acts as a checkpoint and stores an additional information: $Offset$, stored on 64 bits. The $Offset$ of the slot $i$ is the distance between $i$ and the last slot of its run. We store the $Offset$ of the first slot of each block. Because we store an additional 64 bits number every 64 slots, it increases the number of metadata bits per slot to 3. As shown in [1], we can encode the $Offset$ so that it uses 0.125 bits per slot instead of 1. In our implementation, we still made the choice to use 64 bits for every $Offset$, bringing the number of metadata bits to 3 instead of 2.125 for memory alignment reasons. Thanks to the $Offset$ information, it is now possible to count the number $d$ of runs that started before the slot $i$ and after the first slot ($j$) of the same block. Then we jump to the position given by $Offset(j)$ and we find the $d^{th}$ $runend$ from there.

In summary, we compute $runend\_position(i) = Select(runends[Offset(j), i-1], d)$, with $d = Rank(occupieds[j, i], i)$.

#### Side effect of lowering s

When the number of $s$-mers is increasing faster than the number of $k$-mers for the same dataset, there could be the need to double the size of the BQF with $s$-mers requiring $2^{q+1}$ slots, when $k$-mers would have fit in a BQF composed of $2^q$ slots. We created a synthetic dataset for generating such a situation, we used *sea-water34M* numbers of $s$-mers for every value of $s$ and simply added 100 million elements everywhere. This is for visualisation purposes only. Figure 1 shows that, when $s = 16$ or $s = 17$, more than $2^{29}$ elements are to index so $2^{30}$ slots are needed. For these two values of $s$, we can notice that we use even less bits per element ($high\_bound$ and $low\_bound$) than for higher $s$ values, despite the doubling of the structure size. This is because the more elements we add, the more efficient the structure becomes. This being said, in practice, doubling the size of the structure means that the load factor drops from 95% to 50%, *i.e.* space efficiency at that moment jumps from $low\_bound$ to $high\_bound$. To sum up, in some cases, decreasing $s$ might have a momentary negative impact but the overall space efficiency of the BQF continues to improve. It is always beneficial to decrease $s$ until we reach the construction false positive threshold, $s = 17$.

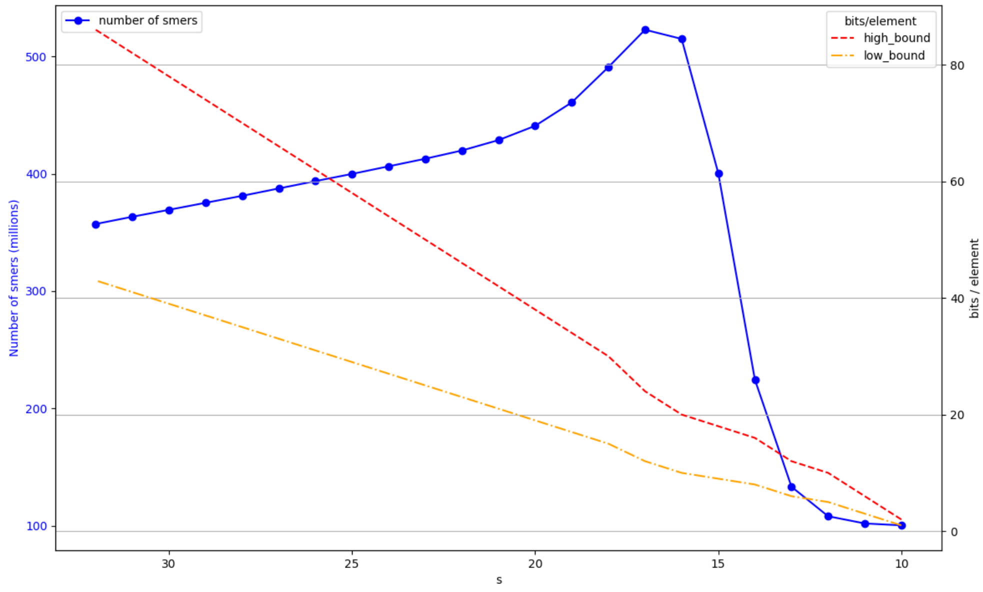


Figure 1: Effect of $s$ on the number of $s$-mers in a fictitious dataset (plain blue line). The data has been modified from dataset *sea-water34M* so that the number of $s$-mers reaches a threshold: $2^{29}$ (500M), requiring to double the number of slots. Space efficiency is also plotted (dashed lines) in bits per element with both boundaries: $high\_bound = 50\%$ load factor, $low\_bound = 95\%$ load factor.

#### Metagenomics complexity

Figure 2 shows the complexity of metagenomics dataset. Because of the variety of species sequenced in metagenomics samples, a vast majority of $k$-mers are present once. Additionally, 99% of $k$-mers are present less than 10 times in *sea-water34M* dataset, and 83% for *gut* dataset. Thus the challenge here is to index low redundant and low abundant $k$-mers.

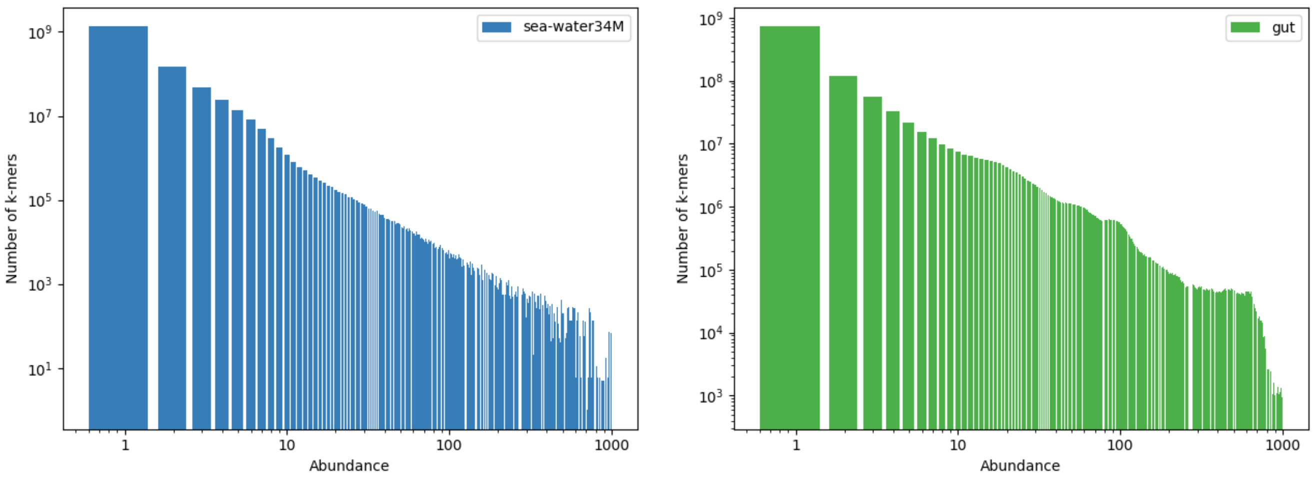


(A) (B)

Figure 2: $k$-mers numbers over (A) *sea-water34M* dataset an (B) *gut* dataset. Abscissa and ordinate are on a logarithmic scale.

### References

[1] Prashant Pandey, Michael A. Bender, Rob Johnson, and Rob Patro. A General-Purpose Counting Filter: Making Every Bit Count. In *Proceedings of the 2017 ACM International Conference on Management of Data*, SIGMOD '17, pages 775–787, New York, NY, USA, 2017. Association for Computing Machinery.
